# Modeling cancer dependency with deep graph models

**DOI:** 10.1101/2024.02.26.582022

**Authors:** Hengyi Fu, Bojin Zhao, Peng Wang

**Affiliations:** Department of Biomedical Sciences, Faculty of Health Sciences, University of Macau; Ministry of Education Frontiers Science Center for Precision Oncology, University of Macau

## Abstract

A fundamental premise for precision oncology is a catalog of diverse actionable targets that could enable personalized treatment. Large scale Genome-wide lost-of-function screens such as cancer dependency map have systematically identified single gene vulnerabilities in numerous cell lines. However, it remains challenging to scale such analyses to many clinical samples and untangle molecular networks underlying observed vulnerabilities. We developed a deep learning framework, DepGPS, combing graph neural networks with transformers to model the network interactions underlying tumor vulnerabilities. Our model demonstrated an improved ability to predict context-specific vulnerabilities over existing models and showed a higher responsiveness in perturbation analysis. Furthermore, perturbation induced dependency changes by our model demonstrated utility to support context-aware identification of synthetic lethal genes. Overall, our model represents a valuable tool to extend tumor vulnerability analyses to broader range of subjects and could help to decipher molecular networks dictating context-specific tumor vulnerabilities.

## Introduction

Cancer is a genetic disease driven by mutations and currently accounts for one sixth of all deaths worldwide^1^. The mechanism of cancer evolution leads to considerable heterogeneity in patients, such that there are multiple cancer clones driven by different combinations of mutations in the tumor of the same patient^2^. A promising strategy to overcome heterogeneity and cure cancer is personalized targeted therapy. Precision targeted therapy aims to first identify cancer-dependent genes, whose inhibition can cause cancer cells to die or stop growing, and then selectively kill cancer cells with drugs that target specific cancer-dependent genes^3^. The first successful case of personalized targeted therapy is to target BCR-ABL fusion genes for the treatment of chronic myeloid leukemia (CML)^4^, which increased the 5-year survival rate of CML patients to more than 90%.

The underlying hypothesis of precision cancer medicine is that cancers have essential genes (cancer-dependent genes) that can be used to develop less toxic targeted drugs to treat cancer by targeting and inhibiting cancer-dependent genes. The success of precision cancer medicine depends on our ability to identify and treat specific weaknesses in patients’ tumors. Ideally, we would identify all the cancer-dependent targets in human cancers, all the combinations of these targets that sustain cancer and develop drugs to inhibit each target. Finally, we need non-cross-resistant therapeutic combinations to overcome subclone heterogeneity in human tumors. But we are still far from that goal, and less than a quarter of cancer patients currently benefit from precision medicine^5, 6^. There are two root causes for this dilemma: 1) we lack drugs that target most known cancer-dependent genes, and 2) we lack a systematic and in-depth understanding of cancer-dependent genes, resulting in many patients without known targets for treatment.

To identify novel therapeutic targets for precision cancer therapy, a growing number of studies are performing cancer-dependent analyses on a genome-wide scale and on in vitro cancer models^7-15^. Of particular note are two large pan-cancer CRISPR screening conducted by the Broad and Sanger Institutes^7, 16^ The two institutes have also joined forces to publish a report on Cancer Cell Line Encyclopedia (CCLE) cell lines to construct a comprehensive atlas of all intracellular dependencies and vulnerabilities of cancer: the Cancer Dependence Atlas (DepMap). At the same time, DepMap also contains multi-dimensional omics data and drug response data of CCLE cell lines, which is expected to greatly promote precision cancer medicine.

The goal of cancer dependency mapping research is to be able to quickly establish a cancer-dependent gene map of each cancer patient’s sample to guide precision cancer treatment. However, the current cancer dependence atlas cannot meet the requirements of precision cancer treatment. First of all, the results of DepMap’s study show that in addition to mutated genes, there are a large number of genes that are not mutated, but are still necessary for cancer cell growth and survival, and changes in gene expression, copy number changes (CNV), mutations, etc., of the genes themselves and related regulatory factors will make the genes alter or even lose their tumor dependence^8^. Because the gene expression, CNV, and mutation profiles of cancer patients are significantly different from those of cancer cell lines and are highly heterogeneous, many tumor types are underrepresented or completely absent in CCLE cell lines. As a result, the tumor-dependent data obtained by DepMap in cell lines often cannot be directly applied in cancer patients. Second, because the construction of organoids or mouse Xenograft models are costly and time consuming, it is challenging to directly screen dependent genes on patient tumor samples utilizing DepMap’s experimental technologies^17^.

Mining the relationship between gene expression, CNV, mutations, etc., and cancer gene dependencies through machine learning can not only help reveal the mechanism of cancer gene dependencies, but more importantly, also provide an efficient method to establish personalized cancer-dependent gene maps from tumor patient omics data, which is of great value for promoting precision cancer diagnosis and treatment. Currently, majority of the available methods still rely on traditional machine learning methods, and could not generalize well to unseen samples^18-21^ ? Currently only one deep learning method based on autoencoders, deepDep, has been developed to predict cancer dependencies^22^. Although deepDep achieved good prediction on the BROAD CRISPR dataset (test set correlation coefficient 0.87). However, the method did not perform well on RNAi-based validation data (independent validation correlation coefficient 0.47-0.66), suggesting that we need more effective approaches to address this problem, which is of great value for precision cancer treatment.

In recent years, the development of transformer and graph neural networks (GNN) related deep learning methods have achieved a series of amazing successes in different fields such as natural language processing, image generation, biology, etc^23-25^. One of the most notable results is alphafold2^25^. Alphafold2 has introduced attention mechanisms into graph neural networks and has achieved major success in protein structure prediction. The success of alphafold2 shows that if the problem or data has a network structure, then the establishment of a framework combining GNN and transformer will not only be able to use GNN to effectively extract the local features of the network, but also could use the transformer to efficiently learn long range relationships, and the effective combination of the two could provide state-of-the-art solution. The GraphGPS framework which combined GNN with transformer has obtained state-of-the-art results across a diverse set of benchmark datasets, confirming the benefits of this architecture^26^.

The DepMap results have shown that gene expression, CNV changes and mutations determine the dependence of genes in the corresponding tumor through associated regulatory networks^7, 16, 27, 28^. Thus, integrating bio-network information with multi-dimensional omics data may significantly improve the ability of models to predict cancer-dependent genes. Here, we combined transformer with GNN utilizing large-scale protein-protein interaction network (PPI) information to integrate multidimensional omics data to predict cancer-dependent genes. We also leveraged pretrained embedding of genes (gene2vec) and various positional and structural embeddings to further enhance model performance. Independent test confirmed that our model achieved better performance than previous methods. Furthermore, our model better captures sematic interaction among genes and are much more responsive in perturbation analysis than deepDep, and consequently, can identify cell line specific synthetic lethal genes. Finally, we established a map of druggable targets in clinical samples by integrating predicted cancer-dependent genes for the TCGA dataset with drug target information. In summary, we established an efficient method for establishing personalized cancer gene dependence maps, identifying potential regulators of cancer dependency and candidate drug regimens for patients, which will play a positive role in promoting precision cancer research and personalized clinical treatment.

## Results

### Overview of the DepGPS model

The first step of DepGPS learning pipeline is to convert multi-omics data to tokenized representation for learning with the transformer and graph neural network architecture (Fig. 1A). The discrete CNV and mutation profiles are directly tokenized as detailed in materials and method section. For the float gene expression data. We employed a Gaussian Mixing Model approach to estimate the number of clusters and corresponding mean and variance. The float gene expression values were then separated into different bins according to number of standard deviations away from cluster mean. In current implementation, we separated gene expression into 10 bins which corresponding to 10 tokens.

**Figure 1.**
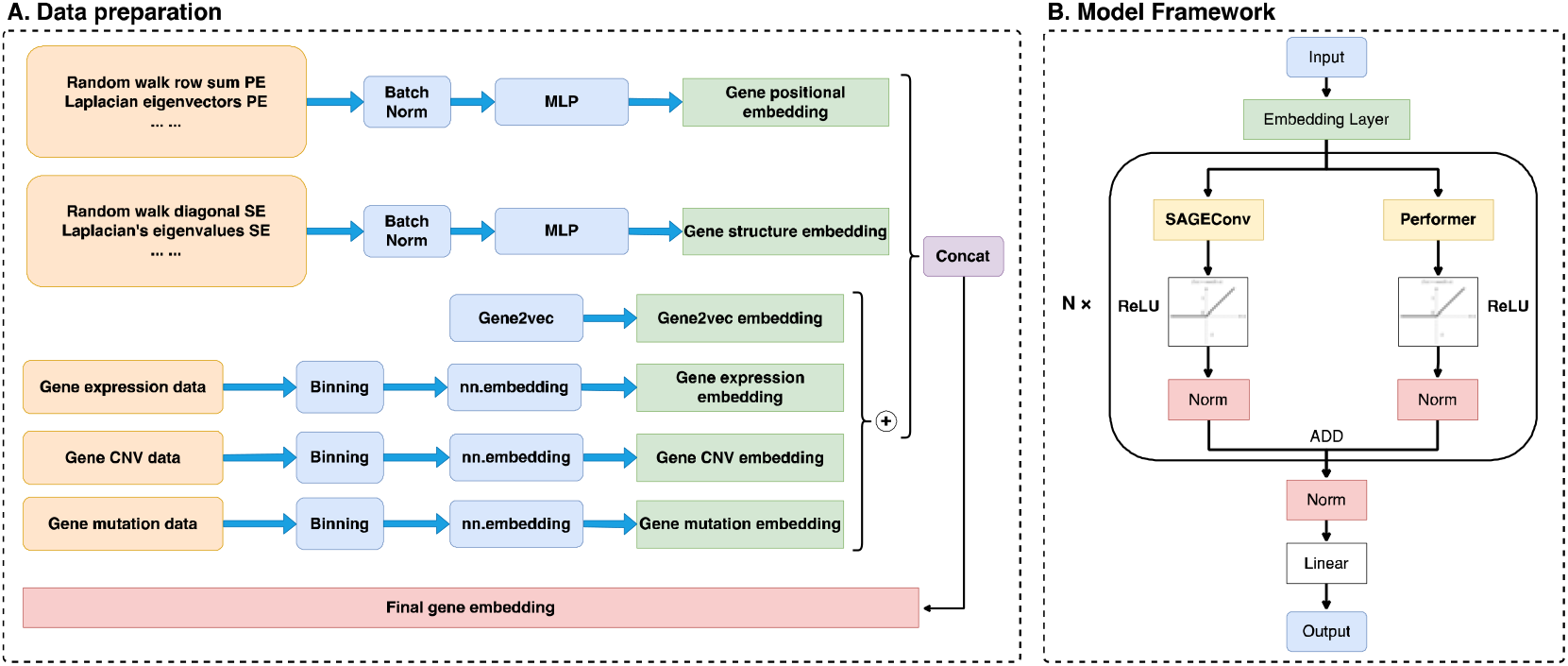
Data preparation pipeline and model architecture of DepGPS. (A) Embedding methods used to convert muti-omics data and topological information contained in PPI into gene embedding. (B) Details of DepGPS architecture.

We also adopted the transfer learning paradigm and utilized gene2vec encoding for genes previously estimated from gene expression data. The gene2vec representation were then add together with the gene expression, mutation and CNV representation to form the finalized gene embedding. To further enhance learning, positional and structural embedding derived from the BioGRID PPI network were concatenated to the gene embedding to generate input embeddings for each gene (Fig. 1A).

The core component of the DepGPS model is a layer consisted of a parallel combination of GNN and Transformer (Fig. 1B). In each layer, embeddings are fed into GNN and transformer separately, and outputs of GNN and transformer are added together to generate output for the basic layer. In our current implementation, the basic layer is repeated two times before the output were forwarded to the prediction layer. Finally, we used a MLP to generate the final predicted dependency scores from the gene embeddings learned by the combination of GNN and transformer.

### DepGPS accurately predicts context-specific cancer gene dependencies

Currently there are an array of different variants of GNN and transformers. We explored several combinations in our model and find that the combination of GraphSAGE, which employs computational graph sampling, and performer, a linear version of transformer, delivered excellent performance and scalability (Fig. 2 and 3). For the combination of GraphSAGE and performer, we achieved a best train Pearson correlation coefficient of 0.94 for the training data and a Pearson correlation coefficient of 0.91 on testing data. We generally achieved best performance within 500 epochs, suggesting that the model is efficient. We observed spikes in the train curve which is consistent with documented behavior of transformers. Furthermore, the test performance of the GraphSAGE and performer combination is close to corresponding training performance (0.94 v.s. 0.91), suggesting that the model could generalize very well (Fig. 2A).

**Figure 2.**
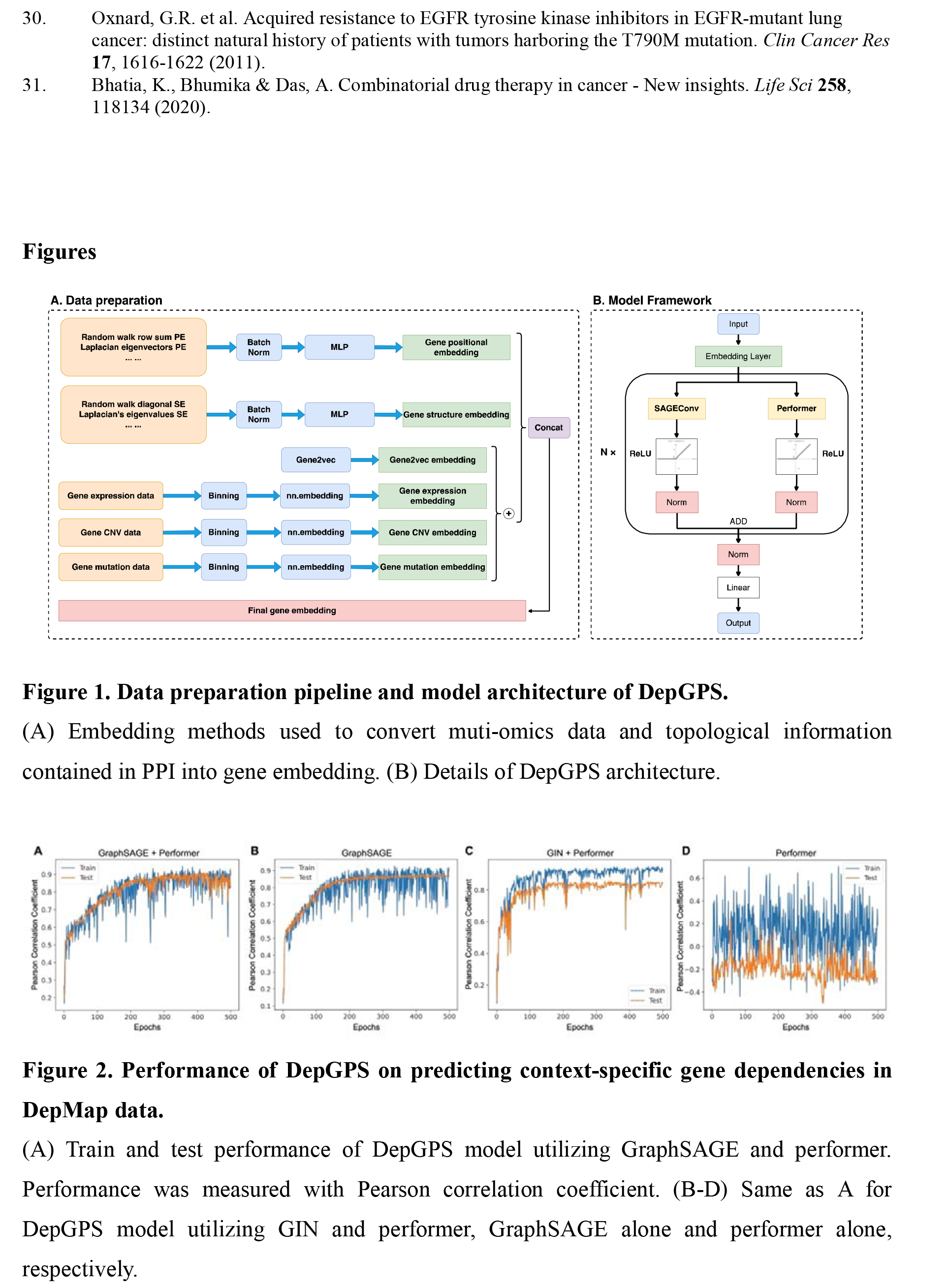
Performance of DepGPS on predicting context-specific gene dependencies in DepMap data. (A) Train and test performance of DepGPS model utilizing GraphSAGE and performer. Performance was measured with Pearson correlation coefficient. (B-D) Same as A for DepGPS model utilizing GIN and performer, GraphSAGE alone and performer alone, respectively.

**Figure 3.**
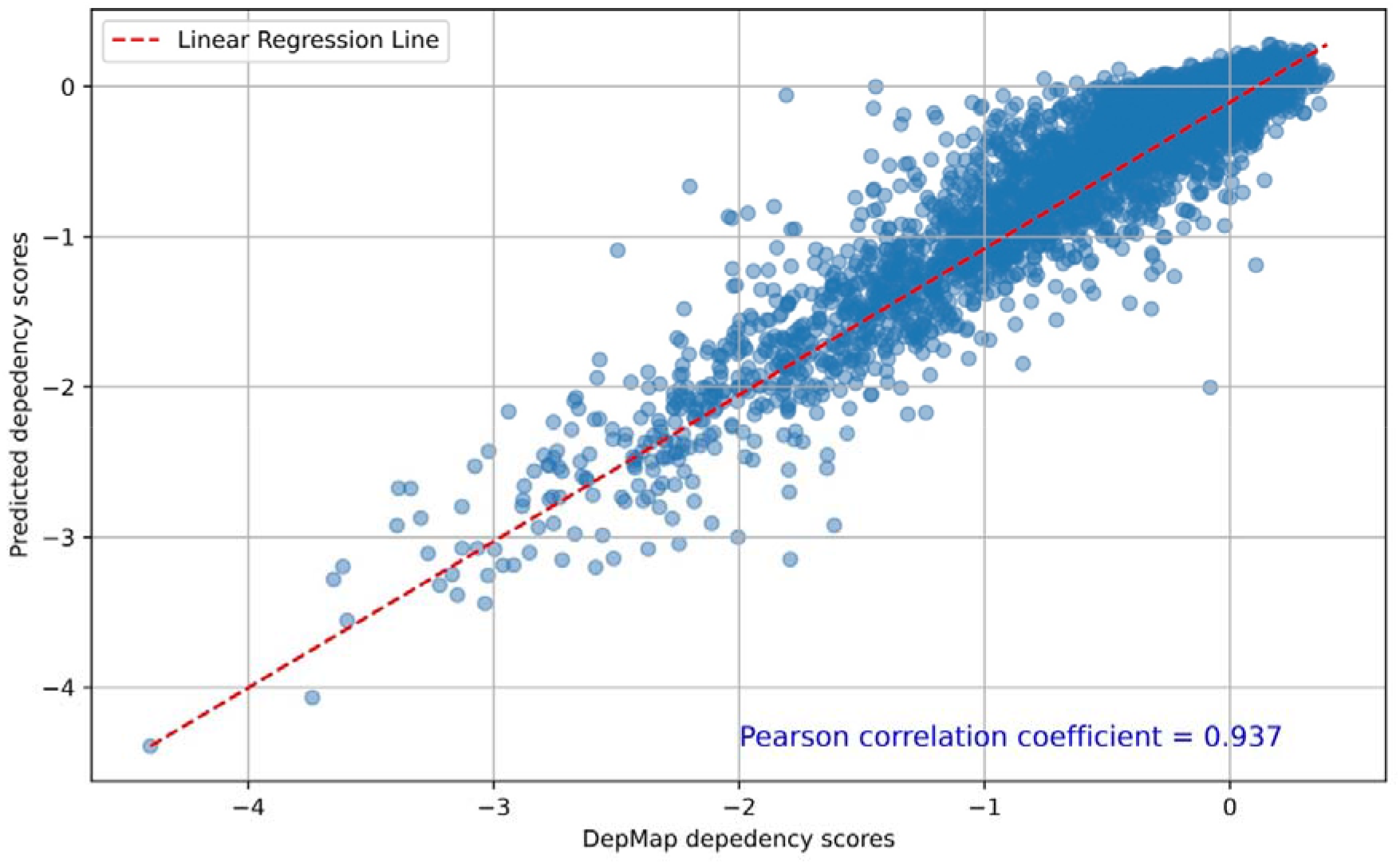
Predicted gene dependency scores demonstrate high correlation with experimentally measure gene dependencies. Scatter plots showing the predicted gene dependency scores v.s. gene dependency scores produced by DepMap project for DMS53 cell line.

We also tested the combination of GIN, which is a more powerful GNN variant with performer. This combination converged faster than the GraphSAGE and performer combination and could achieve a Pearson correlation coefficient of 0.9 within 100 epochs of training (Fig. 2B). The training performance of GIN and performer combination also outperformed the GraphSAGE and performer combination and can achieve a best training Pearson correlation coefficient of 0.95. However, the performance of GIN and performer combination drop to 0.85 for the testing data, suggesting that the GIN and performer combination is less effective in generalizing to new data comparing to the GraphSAGE and performer combination. This observation suggesting that the more powerful GIN variant may lead to overfitting when combined with performer.

Because transformer and GNN alone are powerful architectures that can achieve state-of-the-art performance in many tasks, we also tested the capacity of GraphSAGE or performer to predict dependency scores from multi-omics data. When utilizing the same set of model specifications and training parameters, GraphSAGE alone performed competitively achieving a training Pearson correlation coefficient of 0.92 and a testing Pearson correlation coefficient of 0.87. On the contrary, performer alone could only achieve a training Pearson correlation coefficient of 0.70 and a testing Pearson correlation coefficient of 0.25 (Fig. 2C and D). These results suggested that for the gene dependency prediction task, the local information captured by GNN dictates the model performance while the global syntax captured by transformer provides important details to improve both the predictive performance and generalizability.

### Perturbation analyses with DepGPS identify candidate synthetic lethal genes

Previous analyses of DepMap data suggested that the observed gene dependencies depend on cellular context. Because DepGPS could accurately predict the cell line specific gene dependency scores, we examined whether the DepGPS model could capture the cellular context responsible for observed gene dependency. We adopted a simple perturbation approach similar to deepDep to analyze the context of observed gene dependency (see materials and methods). We first in silico changed a gene’s status (gene expression, mutation and CNV), and calculated the induced changes in all gene’s dependency scores as predicted by DepGPS, where a gene with a negative change indicates that the gene became more essential after the target gene was in silico perturbed.

In sharp contrast to deepDep, where vast majority of induced changes in gene dependency scores is < 0.005 after a gene was perturbed (5th to 95th percentiles, −0.0055 to 0.0054), perturbation of a gene’s status induced changes in gene’s dependency score with a greater magnitude in DepGPS. For example, in A549 cell lines, when KRAS was perturbed, the 5th to 95th percentiles changes in other gene’s dependency scores are -0.0201 to 0.0087, about 4-fold larger that of deepDep (Fig. 4A and B). Interestingly, the perturbation in gene expression induced a much bigger change than these induced by perturbing mutation or CNV status (5th to 95th percentiles -0.0085 to 0.0036, -0.0001 to 0.0013 and -0.0005 to 0.0014 for gene expression, mutation and CNV, respectively) (Fig. 4A). This result is consistent with published analysis results of the original DepMap paper, where they found that gene expression changes could explain majority of context specific gene dependency scores.

**Figure 4.**
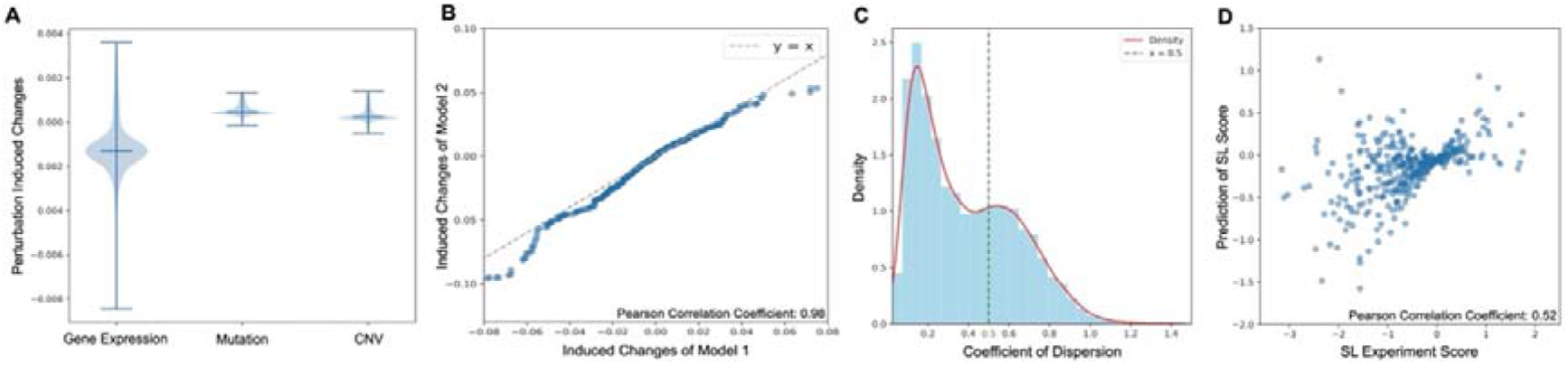
Perturbation analyses of DepGPS. (A) Violin plots showing the distribution of perturbation induced score changes for gene expression, mutation and CNV, respectively. (B) Scatter plot showing high consistency in perturbation induced score changes between different DepGPS models. (C) Histogram showing the distribution of coefficient of dispersion for all genes. (D) Scatter plots showing the predicted SL scores v.s. experimentally determined SL scores for KRAS SL genes in A549 cell line.

The large changes in gene dependency scores indicating that perturbation analysis with DepGPS may help to illustrate mechanisms of observed gene dependencies. Because gradient descent will only produce local optimal results, we first investigated whether the perturbation analysis could generate consistent results across DepGPS models optimized in independent runs. Reassuringly, perturbation induced changes in gene dependency scores are highly correlated between independently optimized DepGPS models, demonstrating Pearson correlation coefficient greater than 0.9 (Fig. 4B). Furthermore, we calculated the coefficient of dispersion of perturbation induced gene dependency score changes for each gene. Expectedly, 99% of genes demonstrated a coefficient of dispersion < 1, confirming that the perturbation analyses results generated by independent DepGPS models are highly consistent (Fig. 4C).

Synthetic lethality (SL), where knocking out a non-essential gene in the presence of specific mutations could causes increased cell death, is a promising approach to identify novel drug targets. Because gene dependency could be modulated by the status of other genes, we explored the ability of DepGPS model to infer SL genes. KRAS is a prominent oncogene and is generally considered undruggable. Consequently, several genome-scale screening experiments have been carried out to identify KRAS SL partners.

SL is a complex phenomenon that demonstrates significant context dependency. Hence, the capacity to identify SL genes in a cell line specific context-aware manner will be of great interest both for illustrating mechanisms of SL and the development of personalized target therapy. We investigated the ability of perturbation analysis to identify context-aware SL genes. To this aim, we utilized 20 trained DepGPS models requiring the model achieves a testing Pearson correlation coefficient > 0.9. We then systematically perturbed KRAS in A549 cell line and calculated the induced changes in gene dependency scores at genome scale. We then designed a simple supervised strategy to integrate the changes of genes’ dependency score across all 20 models by utilizing a simple two-layer neural network. The network was trained on genes scores generated by CRISPR-CAS9 double knockout screening performed in A549 cell line and predicts a final SL score for each gene. This simple integration procedure achieved a Pearson correlation coefficient of 0.52 between predicted SL gene scores and experimentally determined SL scores in A549 cell line (Fig. 4D). These results suggested that the perturbation analysis of DepGPS model is effective in identifying context-aware SL genes.

### Establishing a map of druggable targets for clinical samples

A bottleneck of precision oncology is the lack of actionable targets to such an extent that about 75% of patients could not benefit from targeted therapy. We next investigated whether the candidate cancer dependent genes predicted by DepGPS in clinical samples could help to alleviate this situation. We first applied DepGPS and predicted gene dependency scores for TCGA lung cancer samples. To establish the set of actionable targets in clinical samples, we first identified potential cancer dependent genes by requiring the candidate genes have a predicted dependency score less than -0.7 and are not housekeeping genes. Finally, we intersected candidate cancer dependent genes with 8 drug-target database and assembled a set of 312 known drug-target pairs containing 41 actionable target genes (Figure 5A).

**Figure 5.**
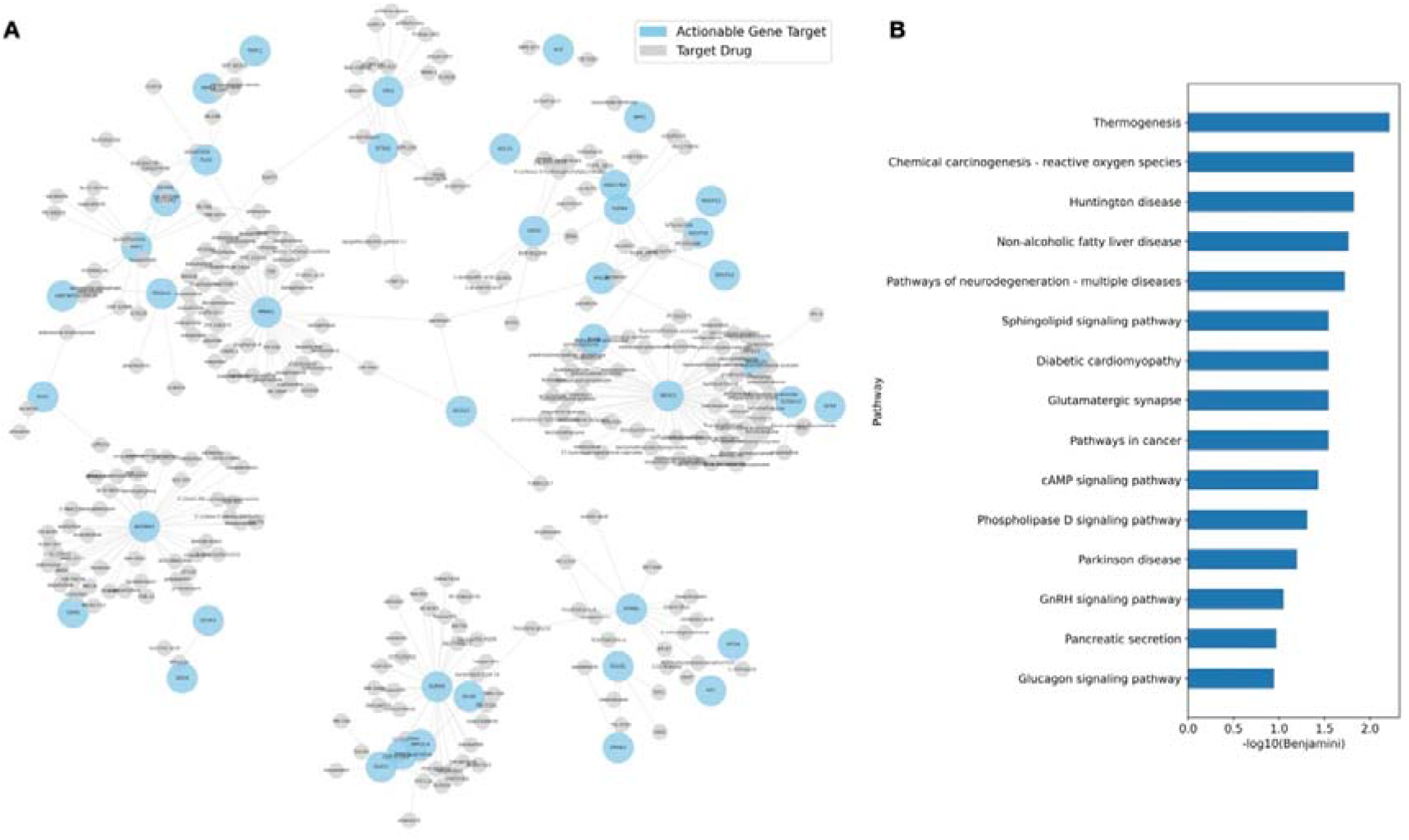
Gene dependency analyses expand scope of actionable targets in clinical samples. (A) Network showing actionable genes identified from predicted dependency scores and corresponding drugs in TCGA lung cancer. (B) Pathway enrichment results for actionable genes in A.

In contrast to actionable targets derived from mutation/CNV alone, where only 29% of samples contain at least 2 actionable targets, more than 69% of analyzed samples contain >=20 actionable targets based on predicted dependency scores. These putative targets could serve as valuable candidates for further investigation that could substantially expand the impact of precision oncology. Interestingly, the putative actionable targets display a diverse distribution in different pathways (Fig. 5). Because it is less likely to develop cross resistance for targets performing different biological functions, these candidates identified by dependency score could serve as valuable candidates to develop combinatorial therapies.

## Discussion

Personalized targeted therapy holds great promise to cure cancer^29^. Unfortunately, currently over 75% of cancer patients lack actionable targets and won’t benefit from targeted therapy^5, 6^. Moreover, virtually all targeted therapies are plagued by acquired resistance and only short-term benefits are sustained^30^. Combinatorial therapies have been proposed to overcome acquired resistance, which requires numerous targets^31^. Consequently, to expand targeted therapy to more cancer patients, and to develop combinatorial therapies to overcome acquired resistance, we need methods to acquire more actionable targets. Efforts like DepMap have demonstrated great promise to expand the list of actionable targets. However, extending these technologies to large cohort of novel samples remains challenging.

Computational strategies offer a cost-effective alternative to identify putative cancer dependency genes. Previously, deepDep demonstrated the capacity of deep learning models to accurately predict gene dependencies in a context-aware manner. Consistent with the ability of deep learning models to capture sematic relationships, perturbation analyses with deepDep could identify well known SL genes. However, the capacity of deepDep to identify SL genes is constrained by the limited changes generated by perturbation analysis, possibly because deepDep only learned cell level embeddings. By learning gene level embedding, our DepGPS model registered much greater changes in gene dependency scores in perturbation analyses and demonstrated improved capacity to identify candidate SL genes. In combination with known drug target information, we demonstrated that the putative SL genes identified by DepGPS represent candidates for clinical actions. Future experimental validation of candidate drug targets and mechanistic studies may help to extend precision oncology to a broader range of patients.

A fundamental limitation of current perturbation analyses is that only a single gene was perturbed. Biological functions are typically regulated by molecular complex or pathways. Although single gene perturbation could reveal simple regulatory relationships, a deeper understanding calls for network level analyses. Strategies such as Monte Carlo tree search has been exploited to identify subgraphs that could explain a GNN model. However, such generic strategies didn’t consider biological insights and demonstrated limited capacity to identify biologically meaningful regulatory relationships (data not shown). Future studies to develop biology inspired network level perturbation analyses strategies could be essential to fully realize the potential of deep learning model to reveal regulatory relationships underlying modeled biological phenomena.

## Materials and methods

### Data sets

The latest CCLE data (2023Q2) was downloaded from the DepMap website (https://depmap.org) and includes gene expression, CNV and mutation data for approximately 2,000 tumor cell lines. TCGA data was downloaded from cancer.gov (https://www.cancer.gov/ccg/research/genome-sequencing/tcga) and includes gene expression, CNV, mutation, and clinical data for ~11,000 samples from 33 cancers. Among them, the gene expression data of CCLE cell lines and TCGA samples have been integrated with Celligner algorithm to ensure that the model based on CCLE data can effectively predict cancer-dependent genes and related regulators on TCGA data.

The CRISPR-Cas9 loss-of-function screening data comes from the Cancer Dependency Atlas project (DepMap, 2023Q2, https://depmap.org), which contains dependency data on 17,386 genes in 1,086 cancer cell lines, which were processed using the Chronos algorithm. Cancer-dependent data for RNAi screening came from Broad’s Achilles Project (consisting of 17,098 genes-dependent data from 501 cancer cell lines), Novartis’ DRIVE Project (consisting of 7,837 genes-dependent data from 398 cancer cell lines), and the study by Marcotte and colleagues (consisting of 16,056 gene-dependent data from 77 breast cancer cell lines). All RNAi data were reprocessed using the DEMETER2 algorithm and have been downloaded from the link below (https://figshare.com/articles/dataset/DEMETER2_data/6025238).

### DepGPS model

The DepGPS model consistets of three modules: the embedding module; the representation learning module; and the prediction module.

For the embedding module (Figure 1A), we first tokenize the multidimensional omics data. Mutations were represented by 2 tokens to represent mutated or unmutated states. For CNV, we used 5 tokens to represent deep deletion, shallow deletion, diploid, low-level gain, and high-level amplification. We used binning to convert floating-point gene expression data into tokens. We tested two approaches. The first one is based on quantile binning, this method guarantees a similar number of genes in each bracket. The second method is based on how many numbers of variances are the genes away from the sample mean, which ensures that genes in the same bin have a similar probability. We also test different number of bins from 5 to 10. Finally, we used cross-validation to determine the optimal binning method and the optimal number of bins. We also employed gene embedding from gene2vec to further enhance the node features (https://github.com/david-r-cox/Gene2vec). Finally, we used a variety of positional embedding and structural embedding methods in the Pytorch Geometric library (based on Laplacian, random walk, etc.) to establish positional embedding and structural embedding of genes in PPI networks. Finally, the positional embeddings and structural embeddings were combines with the gene embeddings and token embeddings representing gene expression, CNV, and mutation data to characterize each node (gene) on the PPI network. We tested and compared two methods of combined embedding (point-wise addition or concatenation) in the model, and finally choose the optimal combination.

For representation learning module (Figure 1B), we used pytorch-geometric and performer to build GNN and transformer encoder modules to learn the representation of genes. In the GNN module, we tried a variety of GNN models including GraphSAGE, GAT, and GINE. We also tested several transformer models such as performer and BigBird. In our model, each layer of the learning module contains a GNN and a transformer encoder, and the GNN and the transformer encoder process the data in parallel, and the results of the processes are added, and each layer of the learning module also includes an MLP to process the combined data of the GNN and the transformer encoder. Learning modules are layered on top of each other. In the prediction module, we used MLP to predict the dependence of different genes in the tumor based on the representation learned by GNN and transformer.

For model training, we utilized a weighted mean square error or a weighted mean absolute error as the loss function. The purpose of weighting is to give higher weight to tumor-dependent genes. To construct the optimal model, we systematically evaluated different batch sizes, different normalization methods including batchNorm, graphNorm and layerNorm, different GNN isoforms (GAT, GIN and grapgSAGE), combinations of different transformer encoder isoforms, and different model parameters (such as number of layers, number of attention heads, dropout rate, etc.) through cross-validation. The training of the model was carried out using A100 and V100 at computing platforms of the University of Macau.

### Perturbation analysis

To perform perturbation analysis on a target gene, we carried out computational mutations on candidate regulator genes to simultaneously mutate the gene expression, CNV, and mutation characteristics of the target gene. The strategy is to in silico change a characteristic to the opposite status. For example, the expression of a high expression factor is changed to low expression, and the amplified CNV is changed to deep deletion, and a mutation status is change to normal. We fed the changed gene embeddings into a trained model to simulate their effects on gene dependence. We calculated the in-silico mutation induced gene dependency changes and selected the genes with significant changes as candidate regulators of the target gene’s cancer dependency. Cut-offs for the selection of genes with significant changes were calculated by the simulation. Specifically, we randomly selected genes to perform random in silico perturbation, then calculate the changes of dependency score for all genes. We repeated this process for 10,000 times. Because majority of mutations are passenger mutations with no significant biological impact, the changes in dependency scores formed a normal distribution centered around zero. We then obtained a cut-off value of 99.99% from this normal distribution to select genes with significant changes in cancer dependency scores after the target gene is in silico perturbed.

### Model performance evaluation

The KRAS synthetic lethal genes identified by double knockout screening were downloaded from SynLethDB 2.0 (https://synlethdb.sist.shanghaitech.edu.cn/v2). To evaluate DepGPS, we calculated the Pearson correlation coefficient between perturbation induced changes in dependency scores with SL gene scores documented by double knockout CRISPR-CAS9 screening.

### Pathway enrichment and survival analysis

The gene ontology and pathway enrichment analyses were performed using the enrichGO and enrichKEGG functions from clusterProfiler R-package (v 3.16.0). All enrichment analyses were carried out with default parameters.

